# Mechanics Of Ultrasonic Neuromodulation In A Mouse Subject

**DOI:** 10.1101/2021.09.23.461613

**Authors:** Hossein Salahshoor, Hongsun Guo, Mikhail G. Shapiro, Michael Ortiz

**Affiliations:** Division of Engineering and Applied Science, California Institute of Technology, 1200, E. California Blvd., Pasadena, CA 91125; Division of Chemistry and Chemical Engineering, California Institute of Technology, 1200 E. California Blvd., Pasadena, CA 91125, USA

## Abstract

Ultrasound neuromodulation (UNM), where a region in the brain is targeted by focused ultrasound (FUS), which, in turn, causes excitation or inhibition of neural activity, has recently received considerable attention as a promising tool for neuroscience. Despite its great potential, several aspects of UNM are still unknown. An important question pertains to the off-target sensory effects of UNM and their dependence on stimulation frequency. To understand these effects, we have developed a finite-element model of a mouse, including elasticity and viscoelasticity, and used it to interrogate the response of mouse models to focused ultrasound (FUS). We find that, while some degree of focusing and magnification of the signal is achieved within the brain, the induced pressure-wave pattern is complex and delocalized. In addition, we find that the brain is largely insulated, or ‘cloaked’, from shear waves by the cranium and that the shear waves are largely carried away from the skull by the vertebral column, which acts as a waveguide. We find that, as expected, this waveguide mechanism is strongly frequency dependent, which may contribute to the frequency dependence of UNM effects. Our calculations further suggest that off-target skin locations experience displacements and stresses at levels that, while greatly attenuated from the source, could nevertheless induce sensory responses in the subject.

## 1. Introduction

Ultrasound neuromodulation (UNM), whereby a region of the brain is subject to targeted focused ultrasound (FUS), leading to a perturbation of neural activity [1, 2, 3, 4], has recently received significant attention as a promising technique for neuroscience research and potential clinical applications. Despite its potential, the underlying biophysical mechanisms that are responsible for inducing neural activation are not well understood and constitute an active area of research, (cf., e. g., [5, 6]). For instance, experiments have shown that UNM procedures, in addition to targeted neuromodulation, can activate auditory and somatosensory brain circuits or elicit skin sensations in the subject [7, 8, 9], which may dependent on the FUS frequency [10, 3]. A number of fundamental questions are raised by these observations, including the mechanisms responsible for direct and off-target effects, the frequency-dependence of such effects and their spatial distribution. Computational methods can shed useful light on these questions and contribute to elucidating the effect of the US parameters on the underlying mechanisms.

In this work, we carry out a computational study of UNM in a mouse model using an anatomically detailed finite element representation, organ-dependent material properties including elasticity and viscoelasticity and explicit time-domain numerical integration. Mice are widely used in UNM experiments [12, 13, 10, 4]. Conveniently, their relatively small size renders them computationally tractable and enables the study of UNM effects across a range of FUS frequencies. The geometry of the model is taken from the co-registered x-ray CT scan of a male mouse available through the Digimouse project [11]. The model accounts for major organs such as the skeleton, brain, heart, lungs, liver, stomach, spleen, pancreas, kidneys, testes, bladder, muscles and skin, Fig. 1.

**Figure 1.**
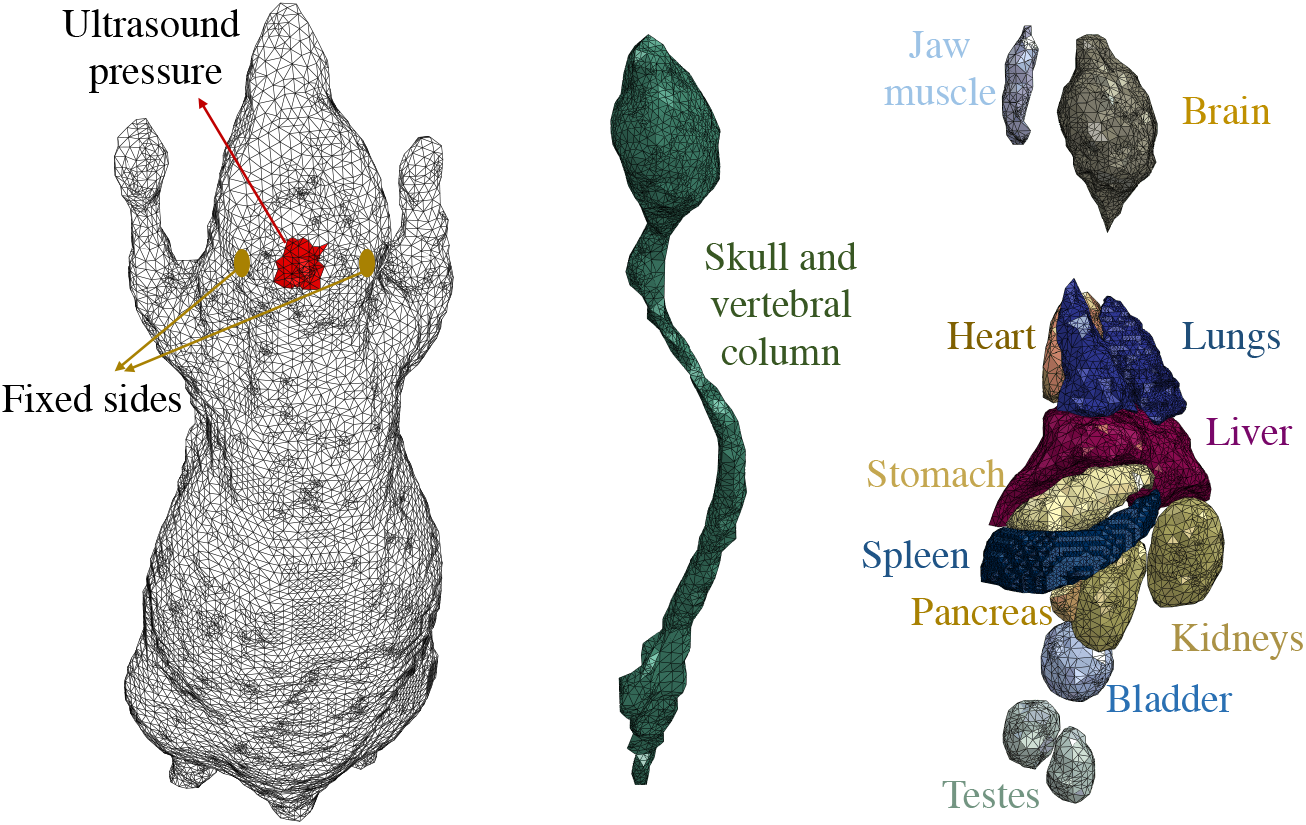
Finite element model of a mouse subject imported from [11]. The left image shows the computational mesh, the location of ultrasound insonation and regions on the skin that are subject to zero displacement boundary condition, replicating a common experimental setup. The middle image shows the cranium and the vertebral column. The right image shows the organs included in the model.

Under the low-intensity FUS conditions of interest here, the induced displacements are of the order of micrometers and the strains are in the permille range. Accordingly, we conduct all calculations in the small-strain linearized-kinematics range. We model the elastic and shear viscoelastic properties of the tissues by means of the standard linear solid model [14] with an exponential relaxation function,

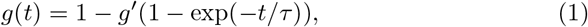

where *g*′ and *τ* are the relaxation coefficient and the characteristic relaxation time, respectively. The material parameters used in calculations for various tissue types in the model are collected from the literature [15, 16, 17, 18, 19, 20] and tabulated in the Table. 1, where *κ* is the bulk modulus, *G* is the shear modulus, and *ρ* is the mass density.

The mouse anatomy, with a total mass of 23.4 g, is discretized into a finite element model with 8.67 million three-dimensional linear tetrahedral elements and 1.54 million nodes, Fig. 1. The element size in the discretization is selected to be less than one tenth of the dilatational wavelength *λ*_*d*_. We recall that *λ*_*d*_ = *c*_*d*_*/ω*, where 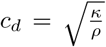 is the dilatational wave speed and *ω* is the frequency of excitation [21]. We simulate insonation by subjecting a region, with an approximate characteristic length of 4 mm, on the skin on top of the cranium to harmonic pressure *P*_0_ sin(2*πωt*) with peak amplitude *P*_0_ = 1 MPa, 1. In order to have the US focused inside the brain, the pressure is imposed in a phased array mode, where a phase lag is introduced in a radial direction. We also prescribe a zero displacement boundary condition at two locations on the skin adjacent to cranium, shown in Fig. 1, which replicates a common experimental setup where bilateral ear bars and/or a head plate are used to fix the mouse head. The calculations are carried out using Abaqus/Explicit (Dassault SystÈmes Simulia, France), which uses explicit time integration with time step of 1.099 × 10^−8^, automatically adopted to ensure the satisfaction of the Courant-Friedrichs-Lewy (CFL) stability criterion [21]. We conducted numerical tests covering a range of mesh sizes and verified that the results of the calculations are sufficiently converged and that numerical dissipation is negligibly small.

We begin by investigating the transmission and focusing of the applied US within the brain. The computed transient pressure wave patterns in a sagittal section of the mouse head through the center of the area of application of FUS are shown in Fig. 2 at four instances of time following the onset of application.

**Figure 2.**
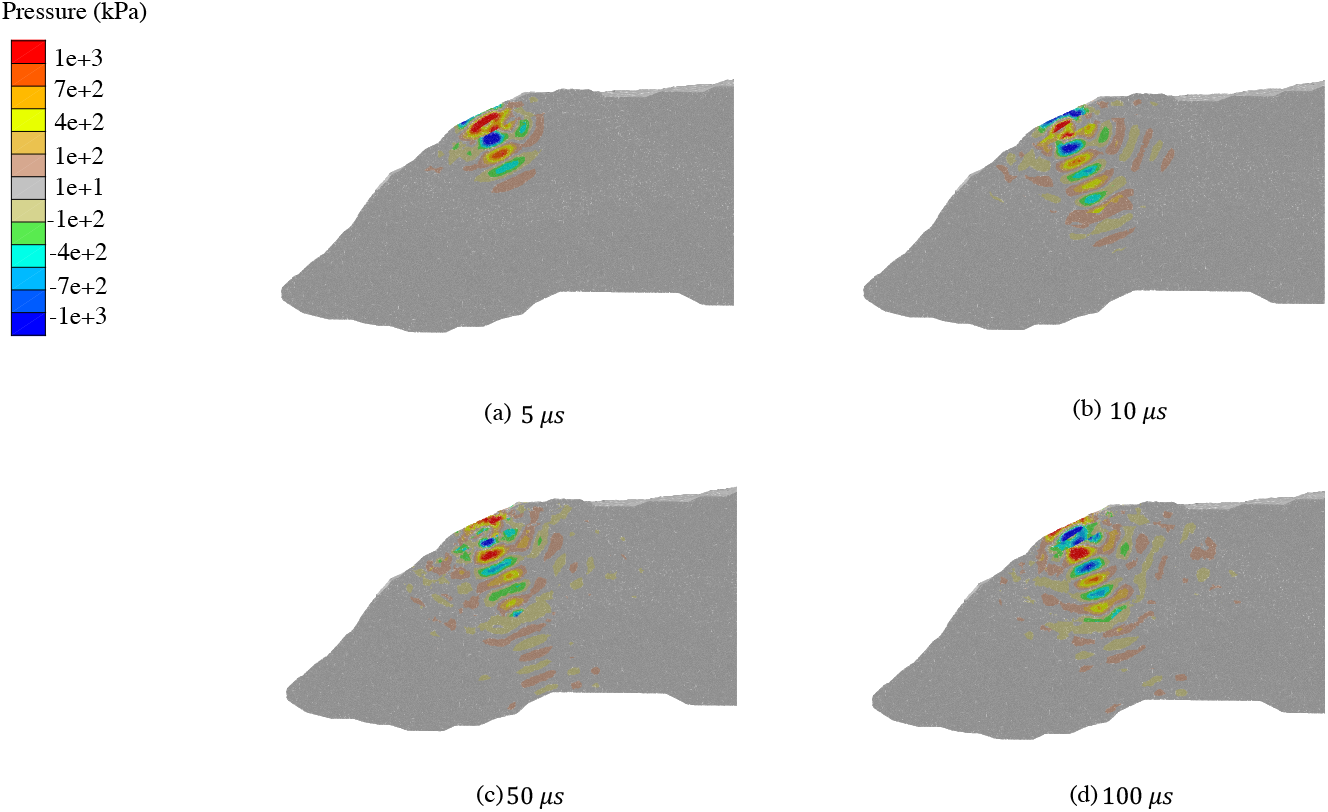
FUS of amplitude 1 MPa and frequency of 400 kHz. Transient pressure wave patterns in a sagittal cross section of the head including the ultrasound focus at: a) 5 *µ*s; b) 10 *µ*s; c) 50 *µ*s; and d) 100 *µ*s.

As may be seen from the figure, the transmitted pressure wave focuses below the cranium and within the brain tissue, reaching an amplitude in the order of 1.5 MPa, indicative of a modest magnification due to focusing. The complex delocalized interference patterns that ensue in addition to the focal signal are also noteworthy. The computed pressure and shear stresses, as measured by the Mises stress, in the focal region over the span of 200 microseconds, corresponding to 80 US cycles, are depicted in Fig. 3. The figure also shows the time history of the maximum principal strain, which remains under 0.2% in amplitude and thus well within the range of small-strain kinematics.

**Figure 3.**
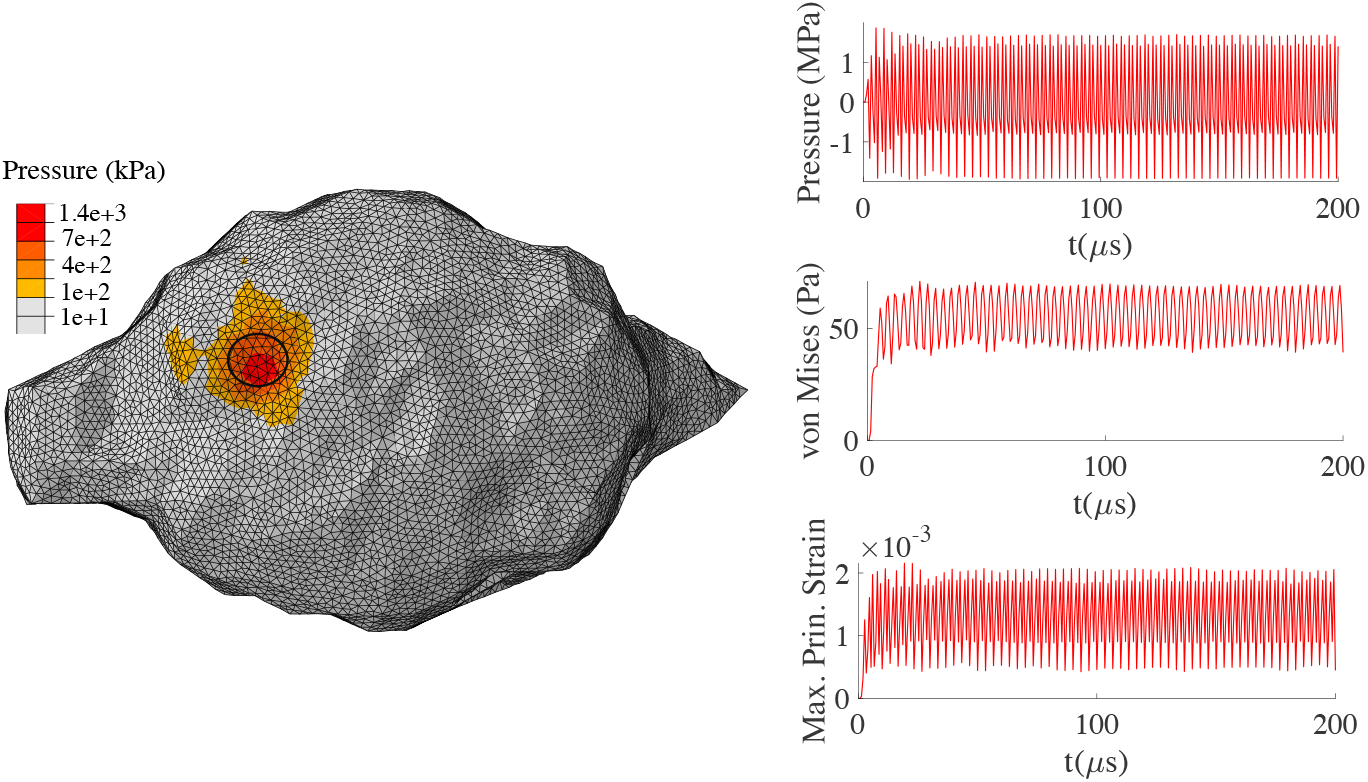
FUS of amplitude 1 MPa and frequency of 400 kHz. Left: Pressure contours on the surface of the mouse brain after 20 *µ*s of insonation. Right: Time history of pressure, Mises stress and maximum principal strain induced in at the focal region (circle) inside the brain.

Remarkably, the shear stresses attain very low values of the order of 50 Pa within the brain. These low values evince the insulating effect of the cranium, which effectively carries the shear waves away from the head into the vertebral column, as well as the low energy/momentum absorption of the US by the soft tissue. A similar shear-wave insulating effect of the brain by the cranium, or ‘cloaking’, is observed in simulations of FUS applied to the human head [22]. The transmission coefficients for waves crossing the various soft tissue and bone interfaces shed quantitative light on the extreme pressure-shear asymmetry observed in the calculations. The values of the transmission coefficients computed from the material properties in Table 1 are collected in Table 2 (cf., e. g., [21] for analytical expressions). We see from this table that, whereas pressure waves cross the interfaces relatively unimpeded in either direction, the transmission of shear waves exhibits extreme asymmetry depending on the direction of propagation, with extremely poor transmission from bone to soft-tissue. This poor transmissibility in turn explains the low ability of the elastically-stiff cranium to transmit shear waves to the elastically-soft brain and the low level of shear stress that ensues in the brain thereof.

**Table 1:**
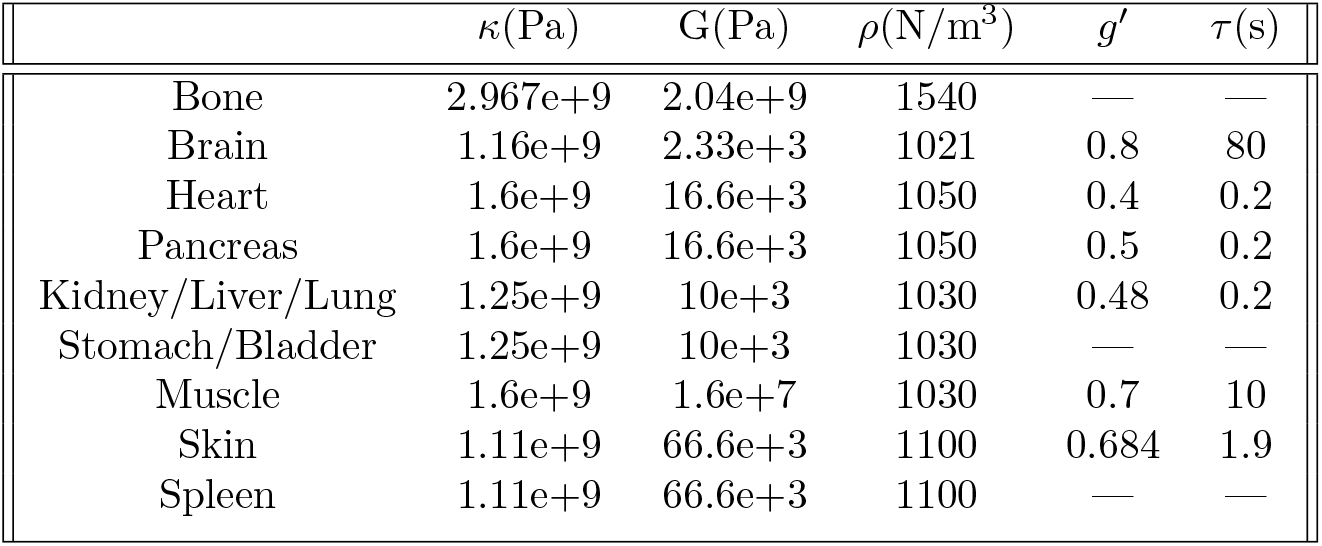
Material properties of different organs in the mouse model [15, 16, 17, 18, 19, 20]

| | $\kappa(\text{Pa})$ | $G(\text{Pa})$ | $\rho(\text{N/m}^3)$ | $g'$ | $\tau(\text{s})$ |
| --- | --- | --- | --- | --- | --- |
| Bone | 2.967e+9 | 2.04e+9 | 1540 | — | — |
| Brain | 1.16e+9 | 2.33e+3 | 1021 | 0.8 | 80 |
| Heart | 1.6e+9 | 16.6e+3 | 1050 | 0.4 | 0.2 |
| Pancreas | 1.6e+9 | 16.6e+3 | 1050 | 0.5 | 0.2 |
| Kidney/Liver/Lung | 1.25e+9 | 10e+3 | 1030 | 0.48 | 0.2 |
| Stomach/Bladder | 1.25e+9 | 10e+3 | 1030 | — | — |
| Muscle | 1.6e+9 | 1.6e+7 | 1030 | 0.7 | 10 |
| Skin | 1.11e+9 | 66.6e+3 | 1100 | 0.684 | 1.9 |
| Spleen | 1.11e+9 | 66.6e+3 | 1100 | — | — |

**Table 2:** Transmission coefficients at the skin-bone interface for both transverse and longitudinal waves reaching the interface through either skin or bone

| Medium of the incident wave | Transverse wave | Longitudinal wave |
| --- | --- | --- |
| Bone | 0.0096 | 0.6811 |
| Skin | 1.99 | 1.31 |

Next, we focus on the role of the vertebral column as a waveguide that carries waves away from the head of the mouse (cf., e. g., [23, 21]). Fig. 4 shows contours of Mises stress in the skull and the vertebral column (spine) at four different instances of time. We observe that after 50 *µ*s the waves are carried away towards the tail, evincing the role of waveguide played by the skeleton and, in particular, the vertebral column. This waveguide effect owes to the large compliance contrast between bone and soft tissue and results in the propagation of waves to locations well outside the US focus. Fig. 5 further depicts the history of Mises stress for FUS frequencies 200, 400 and 600 kHz at four locations of the cranium/vertebral column system: i) the cranial region above the point of application of the ultrasound; ii) the cervical vertebrae (neck); iii) the lumbar vertebrae; and iv) the caudal vertebrae (tail). We observe considerable attenuation of the waves as they travel from the skull to the caudal vertebrae, resulting in a two order-of-magnitude reduction in amplitude. Strong attenuation occurs in waveguides embedded in soft media as a result of leak-off to a degree that is strongly frequency and system dependent. Indeed, we observe from Fig. 5 that attenuation increases markedly with frequency, the higher frequency of 600 kHz resulting in much smaller wave amplitudes than the smaller frequency of 200 kHz.

**Figure 4.**
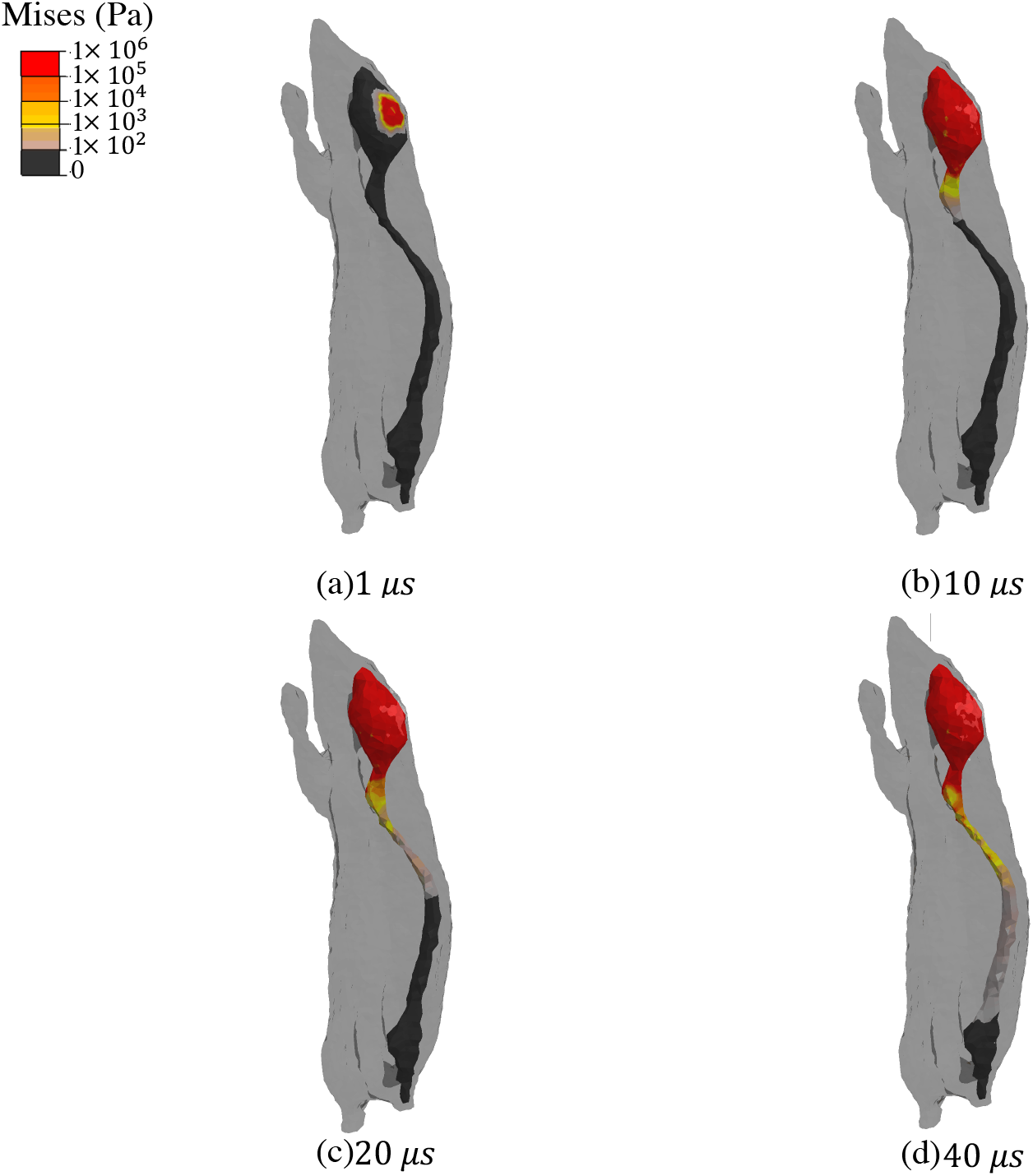
FUS of amplitude 1 MPa and frequency of 400 kHz. Transient shear wave propagation along the vertebral column. Snapshots of Mises stress at: a) 1 *µ*s; b) 10 *µ*s; c) 20 *µ*s; and d) 40 *µ*s.

**Figure 5.**
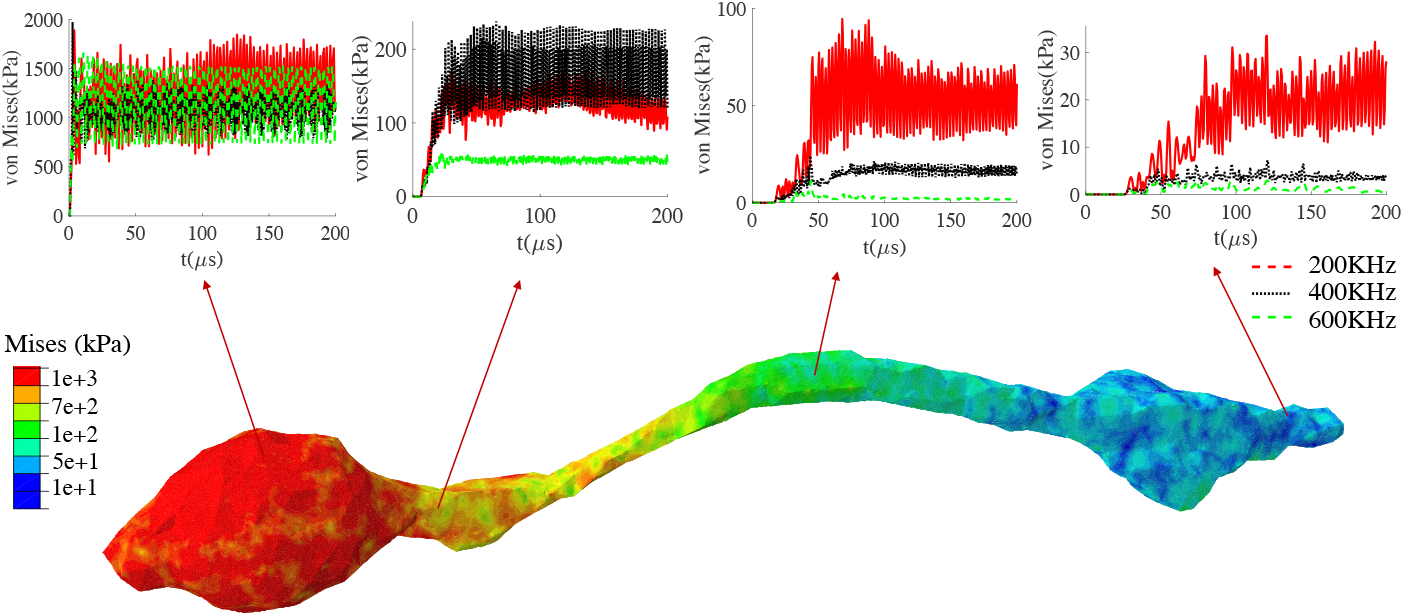
FUS of amplitude 1 MPa. Bottom: Distribution of Mises stress after 200 *µ*s of 400 kHz insonation. Top: Histories of Mises stress at cranial region above the point of application of the ultrasound, the cervical vertebrae (neck), the lumbar vertebrae and the caudal vertebrae (tail).

Finally, we investigate potential mechanical contributions to off-target sensory effects by examining levels of stress at selected locations on the skin. We select two locations, one on the neck and another on the face, Fig. 6, both more than a centimeter away from the region of US application. Calculations are carried out for five FUS frequencies ranging from 200 kHz to 600 kHz, spanning the common range of neuromodulation studies [24, 2, 25, 26, 27, 28, 29]. Fig. 6 displays the computed histories of pressure and shear stresses at both locations. We observe that peak pressure values of the order of 20 kPa are attained at the face location and of 15 kPa at the neck location. Moreover, we observe that the computed magnitude of the pressure does not exhibit significant frequency dependence.

**Figure 6.**
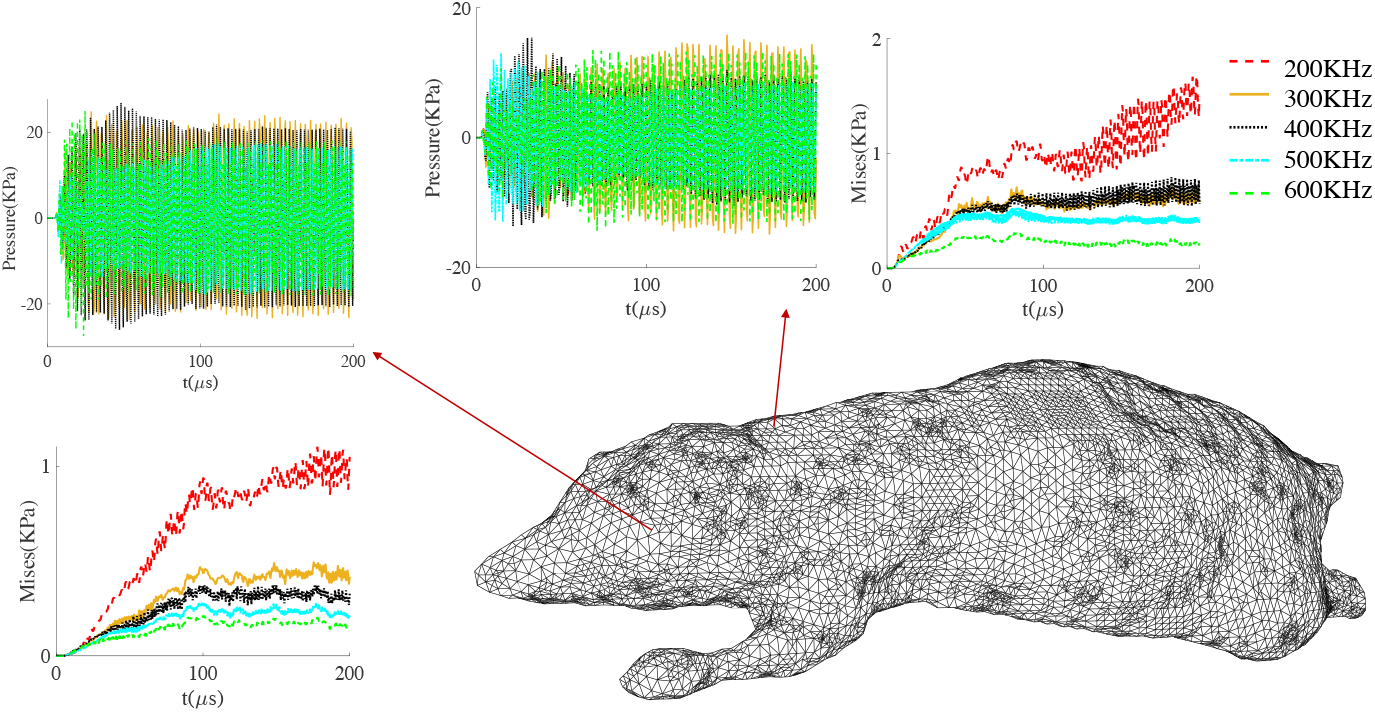
FUS of amplitude 1 MPa. Off-target pressure and Mises stress histories induced at regions on the neck and the face.

In contrast, the computed Mises stresses are significantly smaller in magnitude and exhibit a consistent and marked dependence on frequency, with lower frequencies resulting in larger shear stress amplitudes. The frequency dependence of stress amplitudes could potentially be the result of viscoelasticity. We tested this hypothesis by performing calculations identical in every respect to those shown in Fig. 6 except for turning off viscosity and verified that viscoelastic effects are negligible. We note that the shear wave speed in the skin is 7.8 m/s. Therefore, over the US stimulation time the shear waves recorded on the neck and the face are not transmitted through the skin itself but, instead, carried by the skeleton, which acts as a waveguide. Therefore, the frequency dependence of the shear-wave amplitude shown in Fig. 6 may be attributed to the frequency-dependence of the waveguide mechanism described earlier. By contrast, the pressure waves are carried by all tissues, hence the ostensibly frequency-independent response exhibited by the calculations.

The peak displacements computed at the neck and face are 0.8 *µm* and 0.91 *µm*, respectively. The computed off-target mechanical stresses and deformations, though small, may nevertheless be of sufficient magnitude to elicit a sensory response [30, 31, 32, 33].

In summary, we developed a finite-element model of a mouse, including elasticity and viscoelasticity, and used it to interrogate the response of mouse models to FUS. We find that, while some degree of focusing and magnification of the signal is achieved within the brain, the induced pressure-wave pattern is complex and delocalized, which suggests further research on the optimal control of insonation. In addition, we find that the brain is largely insulated, or ‘cloaked’, from shear waves by the cranium and that the shear waves are largely carried away from the skull by the vertebral column, which acts as a waveguide. Our calculations further suggest that off-target skin locations are excited to levels such as may induce sensory responses in the subject [30, 31, 32, 33]. Our findings also reveal that shear stresses, but not pressures, exhibit a consistent and marked dependence on the FUS frequency, with lower frequencies leading to stronger shear effects, in agreement with observations of UNM effects in rodents [10, 3].

## Acknowledgements

The support of the National Institute of Health through grant 1RF1MH117080 is gratefully acknowledged.

## Notes

### Competing Interest Statement

The authors have declared no competing interest.

